# Loss of PRC2 subunits primes lineage choice during exit of pluripotency

**DOI:** 10.1101/2020.07.08.192997

**Authors:** Chet H Loh, Matteo Perino, Magnus R Bark, Gert Jan C Veenstra

## Abstract

Polycomb Repressive Complex 2 (PRC2) is crucial for the coordinated expression of genes during early embryonic development, catalyzing histone H3 lysine 27 trimethylation. There are two distinct PRC2 complexes, PRC2.1 and PRC2.2, which contain respectively MTF2 and JARID2 in ES cells. Very little is known about the roles of these auxiliary PRC2 subunits during the exit of pluripotency. In this study, we explored their roles in lineage specification and commitment, using single-cell transcriptomics and mouse embryoid bodies derived from *Mtf2* and *Jarid2* null embryonic stem cells (ESCs). We observed that the loss of *Mtf2* resulted in enhanced and faster differentiation towards cell fates from all germ layers, while the *Jarid2* null cells were predominantly directed towards early differentiating precursors and neuro-ectodermal fates. Interestingly, we found that these effects are caused by derepression of developmental regulators that were poised for activation in pluripotent cells and gained H3K4me3 at their promoters in the absence of PRC2 repression. Upon lineage commitment, the differentiation trajectories were relatively similar to those of wild type cells. Together, our results uncovered a major role for MTF2-containing PRC2.1 in balancing poised lineage-specific gene activation, providing a threshold for lineage choice during the exit of pluripotency.

**Highlights:**

1. Enhanced and faster differentiation into all three germ layers in *Mtf2* null embryoid bodies
2. *Jarid2* null cells enriched for early differentiating precursors and neuro-ectodermal cell fates
3. MTF2 is critical for the balance of activation and repression of key developmental regulators
4. PRC2 coordinates lineage choice and execution of the lineage-specific program by thresholding of lineage-priming

## Introduction

During early stages of mammalian development, the epiblast receives both inductive and repressive cues to precisely regulate the exit of pluripotency and the onset of differentiation. These cues, especially signaling morphogens, pattern the embryo, establish the body axes and specify lineages through dynamic and temporal gradients, eventually producing different cell types from the distinct embryonic germ layers (endoderm, mesoderm and ectoderm)^1–3^. To do that efficiently, pluripotent inner cell mass cells, and embryonic stem cells (ESCs) derived from them, employ a multitude of highly conserved genetic mechanisms to repress or activate the genes in a spatially and temporally controlled fashion.

Polycomb Repressive Complex 2 (PRC2) is one of the key transcriptional repressors in ESCs. It methylates lysine 27 of histone H3 (H3K27me3), marking the chromatin for compaction, and repressing target genes together with Polycomb Repressive Complex 1 (PRC1)^4–8^. In ESCs, specific lineage and differentiation genes like *Brachyury* (mesoderm) and *Otx2* (early differentiation intermediates and neural ectoderm) are repressed by PRC2 during pluripotency^9,10^. Additionally, PRC2 represses ectopic expression of lineage-specific genes, which is thought to stabilize lineage commitment. The core components of PRC2 include the catalytic proteins EZH1/2, EED, SUZ12 and RBBP7/4^11,12^. In recent years, a number of accessory subunits have been found at sub-stoichiometric levels^13–17^. This led to a classification of two distinct PRC2 sub-complexes. PRC2.1 consists of the core together with one of the Polycomblike proteins (PHF1, MTF2 or PHF19) and EPOP, whereas PRC2.2 contains the core subunits with JARID2 and/ AEBP2. MTF2 recruits PRC2.1 to unmethylated CpG-rich DNA^14,18^, and its significance in mESCs was underscored by the relatively strong genome-wide reduction of PRC2 binding and H3K27me3 enrichment upon the loss of *Mtf2*^18–21^. In PRC2.2, JARID2 can bind DNA and also nucleosomes through recognition of ubiquitylated histone H2A (H2Aub119), a modification catalyzed by Polycomb Repressive Complex 1 (PRC1)^22,23^.

While much work has been done to unravel the molecular mechanisms of PRC2 recruitment in embryonic stem cells, little is known about how the PRC2 subcomplexes affect the exit of pluripotency. In mouse ESCs, the PRC2 core subunit genes *Suz12, Eed* and *Ezh2* are not required for naïve pluripotency, but they are required for the maintenance of pluripotency in the primed state^24^. This presents a question of how PRC2 is regulating the exit of pluripotency, and more specifically, how it affects lineage choice. Furthermore, *Mtf2* and *Jarid2* mutants further revealed the requirement for accessory subunits during embryonic development, with mutants experiencing embryonic lethality (before E15.5 for *Mtf2* and E18.5 for *Jarid2* mutants)^25,26^.

In this study, we explored how PRC2.1 and PRC2.2 mutant mESCs behaved during differentiation. Using single-cell transcriptomic analyses of mouse embryoid bodies (EBs), we observed that the *Mtf2* mutant cells differentiated faster into all germ layers, while the *Jarid2* mutants were slower and predominantly gave rise to early ectodermal precursors. Intriguingly, we found that MTF2 represses key lineage-specific transcription factors and signaling genes which are inherently poised for activation in wild-type undifferentiated cells. With the loss of *Mtf2*, their transcript levels and promoter H3K4me3 modifications were increased, whereas H3K27me3 was decreased, allowing for rapid induction of lineage specification genes. As shown in directed differentiation experiments, this leads to a stronger induction of lineage-specific gene expression in mesoderm and endoderm progenitors, while still allowing alternative lineages to be repressed.

Together, our results outline a critical role of PRC2.1 (MTF2) in controlling the threshold of lineage gene activation by maintaining repression on key lineage transcription and signaling factors, and therefore modulating the state of promoter bivalency. Furthermore, the single-cell resolution of EB differentiation revealed differences in lineage potency upon the loss of PRC2.1 and PRC2.2, which is linked to their effects on H3K27 methylation and derepression of specific genes, uncovering their roles during the exit of pluripotency.

## Results

### Embryoid bodies of PRC2 mutant ESCs differentiate to cell types across all germ layers

Previously, we and others have observed differences in core PRC2 recruitment between the loss of MTF2 (PRC2.1 mutant) or JARID2 (PRC2.2 mutant) in mESCs^13,21,27^. This potentially affects the repression of key lineage transcription factor genes, for example *Otx2*, where EZH2 binding was dramatically reduced in the *Mtf2* null compared to the *Jarid2* null or wild type cells (Fig. 1a).

**Figure 1.**
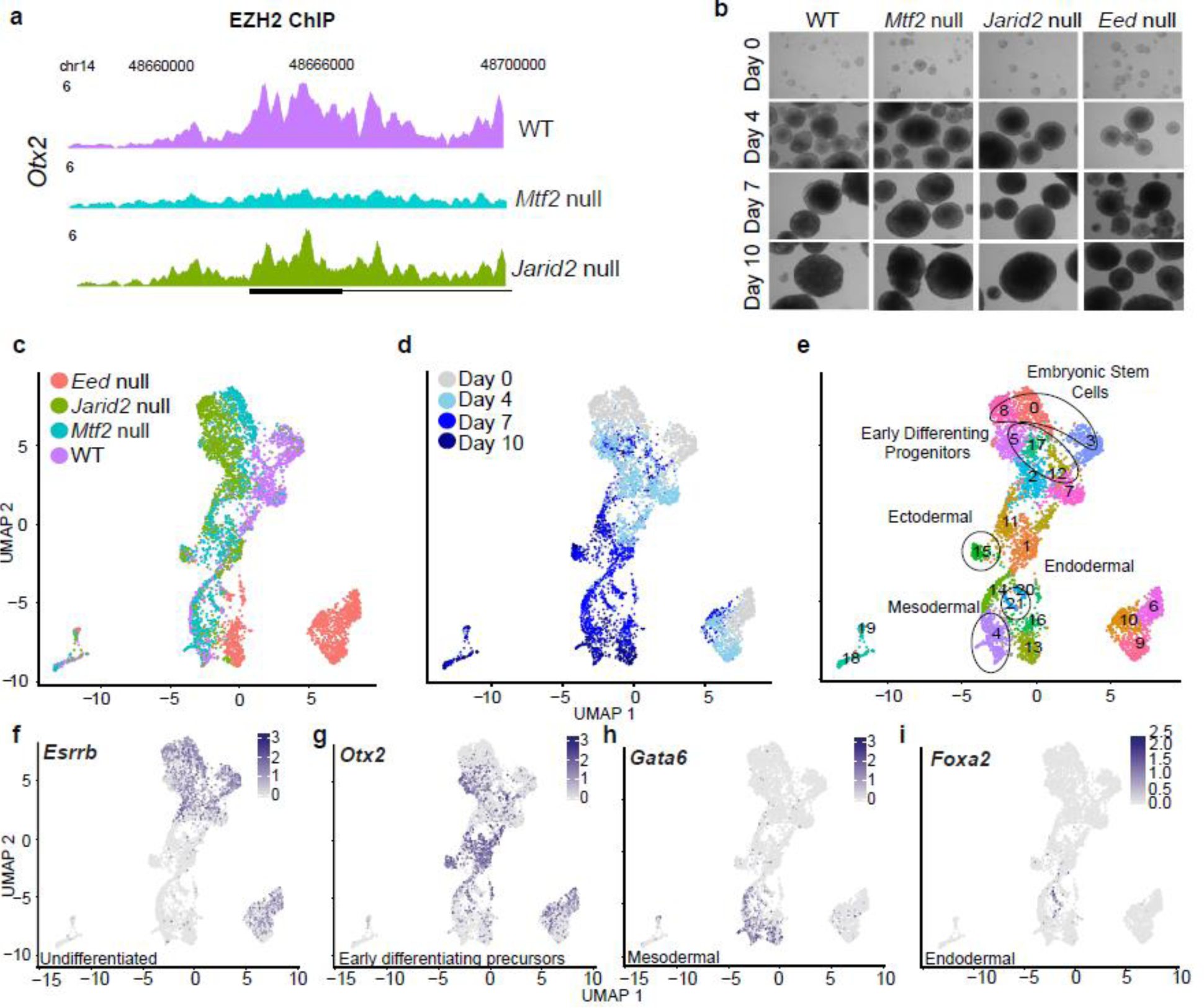
**a**, EZH2 ChIP peak profiles of gene *Otx2* for WT, *Mtf2*GT/GT and *Jarid2*-/- cells. Peaks were RPKM normalized and scaled across backgrounds. **b**, Phase-contrast of Embryoid Bodies from different background and time point of differentiation. **c & d**, Single-cell UMAPs of 4949 cells over all backgrounds and time points. **e**, Single-cell UMAP of cells colored by predicted clusters. **f – i**, Feature maps of selected lineage genes, color intensity based on normalized expression of individual gene.

We wondered how such a loss of PRC2 subunits affects the exit of pluripotency and early germ layer differentiation. To untangle the heterogeneity and track the differentiation lineage potential of these mutants over a broad range of cell fates, we performed mouse embryoid body (EB) differentiation (Fig. 1b) and captured the transcriptomes of close to 5000 single cells encompassing different genetic backgrounds (wild-type, *Mtf2* null, *Jarid2* null and *Eed* null cells) and time points (day 0-10) during differentiation (Fig. 1c-d). Phenotypically, we did not observe any stark differences in shapes and sizes of differentiated EBs between the different genetic backgrounds (Fig. 1b). After stringent quality checks for outliers and drop-outs (cells with little or no mRNA recovered), the remaining cells (n = 4949) were normalized for differences in sequencing depth and batch variation (Methods), clustered using the Louvain method for community detection, and projected in a two-dimensional space using the Uniform Manifold Approximation and Projection (UMAP) method for dimension reduction (Fig. 1c-e).

Clear differences were observed between the different Polycomb mutants during differentiation, particularly between the *Eed* (core PRC2 subunit) null cells and the rest, as expected by the complete lack of PCR2 repression in these cells (Fig. 1c). To confirm this as a molecular phenotype and exclude technical confounders from the SORT-seq protocol, we performed bulk RNAseq on replicated differentiation experiments and observed the same pattern (Fig. 1c-d, Extended Data Fig. 1a). The two-dimensional UMAP representation reproduces the temporal order of differentiation from the expression patterns, with the early and late time points relatively more to the top and to the bottom of the graph respectively (Fig. 1d). The pluripotency markers *Esrrb* and *Nanog* were expressed in cell clusters corresponding to early time points, whereas lineage-specific markers were expressed at later time points (Fig. 1f-i; Extended Data Figure 1c-f). Generally, the undifferentiated ES cells of different genotypes occupy distinct clusters, but some of these cells converge into mixed genotype clusters upon differentiation. This is most dramatically observed for *Eed* null cells, which are quite different from wild type ES cells. Within differentiating EBs, however, some of these cells acquire transcriptomes that are similar to cells with wild type or other genotypes, as observed in clusters 19 and 20 for example (Fig. 1c-d, Extended Data Fig. 1b). Compared to *Eed* null cells, the transcriptomes of *Mtf2* null and *Jarid2* null ES cells exhibit smaller differences with wild type ES cells, but still form genotype-specific clusters when not differentiated (clusters 3, 8 and 0 for respectively wild type, *Mtf2* null and *Jarid2* null cells). In EBs, these undifferentiated cell clusters pass through largely genotype-specific clusters with early differentiation intermediates (respectively 12, 5 and 17), after which they tend to converge into mixed genotype differentiated clusters (Fig. 1c-d, Extended Data Fig. 1b).

To assist a systematic analysis of differentiation potential in relation to the loss of PRC2 subunits, we annotated each of the 22 clusters in our dataset using Anatomy Ontology^28^ and used this in combination with matches to cell types in the recently published Mouse Cell Atlas^29^ (Methods; Extended Data Fig. 2a-b). We then grouped cell clusters into the germ layers for downstream analyses (Extended Data Fig. 2c). The EBs produce a rich diversity of cell lineages of all three germ layers. For example, cluster 4 consisted of cells resembling the lateral mesoderm lineage (e.g. heart and pericardium, mesenchyme, hematopoietic progenitors), expressing high levels of cardiac markers such as *Gata6, Tbx20* and *Isl1* (Fig. 1e, h, Extended Data Fig. 1e, 2a). Clusters 15 and 1 were characterized as ectodermal cells (e.g. neural ectoderm, retina) enriched for *Otx2* expression (Fig. 1e, g, Extended Data 2a). Cluster 21 exhibits endoderm-specific gene expression, including the expression of genes such as *Foxa2* and *Sox17* (Fig. 1e, i, Extended Data Fig. 1f, 2a). Interestingly, many of the clusters appeared to be predominantly enriched for cells from specific PRC2 subunit mutants (Fig. 1c, e, Extended Data Fig. 2d). This raised the question to what extent a bias in differentiation can be caused by differences between the PRC2 mutants starting from the undifferentiated state.

### Altered patterns of lineage commitment in PRC2 mutant embryoid bodies

To assess the patterns of differentiation we used the cell clusters grouped by germ layer and counted the total number of cells from each genetic background and their enrichment over wild type cells (Fig. 2a, Extended Data Fig. 2c-d). *Jarid2* null cells were overrepresented in ectodermal clusters at the expense of mesodermal or endodermal lineages (Fig. 2a). This is mainly on account of *Otx2*-expressing neuro-ectodermal cluster 15, in which both *Jarid2* null and *Mtf2* null cells were overrepresented (Extended Data Fig. 2d). *Jarid2* null cells were also overrepresented in clusters 5 and 17, which represent early differentiation intermediates that also express *Otx2* (Fig. 1e, g). This combination of *Otx2* and pluripotency gene expression, which is also observed in a corresponding cluster of predominantly wild type cells (cluster 12), may correspond to Rosette-stage pluripotency between naïve and primed states^30,31^. Strikingly, cells with a loss of *Mtf2* were overrepresented in all three germ layers, at the expense of clusters with mixed germ layer annotations (Fig. 2a, Extended Data Fig. 2c-d) and early differentiating precursors (negative log 2 fold-change of 2.1 against WT, Extended Data Table 1). The *Eed* null cells, similarly, were enriched in cell types from all germ layers but particularly for endodermal cells.

**Figure 2.**
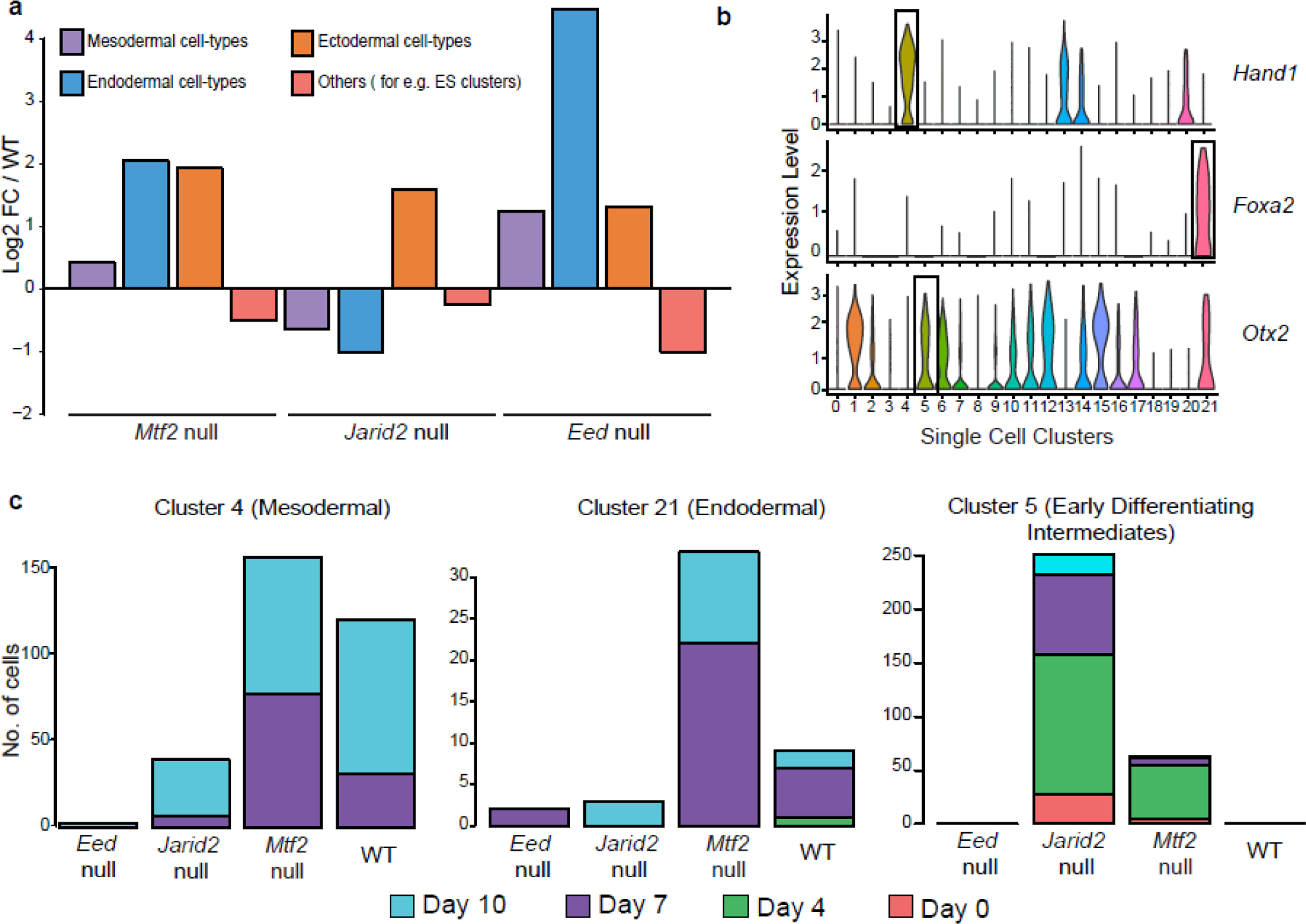
**a**, Barplot showing enrichment of different germ layer cell types from genetic mutants compared against WT. **b**, Violin plots depicting expression of key lineage transcription factors. Boxed out clusters are explored in depth in panel c. **c**, Bar plots of respective lineage clusters showing the number of cells from each time point in each cluster for respective backgrounds.

To understand these enrichments and their relation to time, we examined the cardiac and gut progenitors (respectively cluster 4 and 21) and early differentiation intermediates (cluster 5) in more detail. These clusters express specific transcription factors (Fig. 2b). In cluster 5, one of the clusters with early differentiation intermediates enriched for *Jarid2* null cells, we observed that the cluster contained cells from both late (day 7 and 10) and earlier time points (Fig. 1d, 2c, Extended Data Fig 2e). Similarly, for clusters 4 and 21, which represent mesodermal and endodermal lineages respectively, more *Mtf2* null cells were found in these clusters as compared to wild type and *Jarid2* null cells. This difference is most pronounced at day 7 already, suggesting a rather efficient differentiation of *Mtf2* null cells. Together, these results indicated that the loss of *Mtf2 and Jarid2* affected cell lineage choice upon the exit of pluripotency. *Jarid2* loss was associated with a less efficient transition through differentiation intermediates and with more cells in neuro-ectoderm lineages, while the loss of *Mtf2* resulted in rather efficient differentiation towards lineages of all three germ layers.

### *Mtf2* null cells progress faster through mid-differentiation than WT and *Jarid2* null cells

To observe the process of lineage commitment in more detail, we performed an unsupervised trajectory analysis combining wild type, *Mtf2* null and *Jarid2* null cells. These trajectories provide a common framework to compare the differentiation characteristics of cells with different mutations. Individual cells were ordered based on gene expression differences and plotted as a function of pseudotime (Fig. 3a, Extended Data Fig. 3a-d).

**Figure 3.**
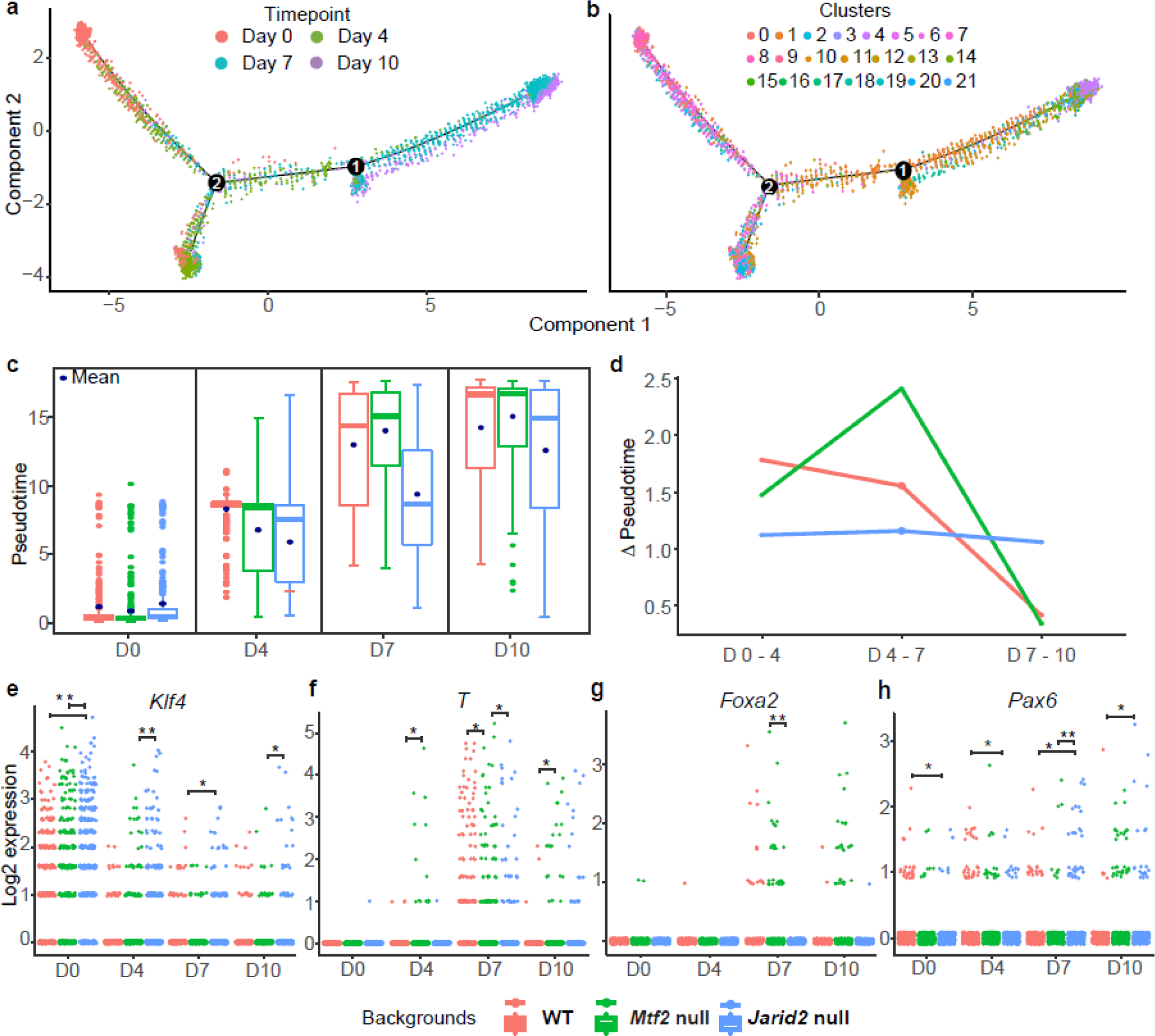
**a**, Trajectory plot of all cells ordered along pseudotime. Two branching points were shown (numbered) along the plot. **b**, Same trajectory plot as (a) but overlaid with cluster identity of individual cell. **c**, Boxplots for each time point showing the pseudotime values of cells from different genetic backgrounds. Dark blue dots represent the mean. **d**, Delta pseudotime plot, showing the distance (change in mean pseudotime value) per unit of time (days) between different genetic backgrounds. **e**, Log2 expression values for selected lineage genes at different time points; comparing between different genetic backgrounds. Each dot represents a single cell. Significant differences (Wilcoxon signed-rank test) are indicated: * p-value < 0.01, ** p-value < 1 × 10^−4^.

By overlaying the experimental time points (day 0, 4, 7 and 10) onto the pseudotime plot, we observed that the differentiation trajectory order of cells reliably recapitulated real-time information (Fig. 3a, b, Extended Data Fig. 3). We observed that *Mtf2* null cells were more concentrated toward the later pseudo-times as compared to the *Jarid2* null cells (Extended Data Fig. 3b-d). To quantify the speed by which the cells differentiated, we calculated the differences in pseudo-time (which is based on gene expression, representing the ‘distance’ of differentiation) over real-time intervals. Strikingly, we found that the *Mtf2* null cells differentiated at a faster rate, especially between days 4 and 7 of differentiation (Fig. 3 c-d). We also found that the *Jarid2* null cells, which were severely delayed by day 7, partially catch up between days 7 and 10 (Fig. 3c).

To relate these differences in trajectory speed to marker gene expression, we examined *T (Brachyury), Foxa2 and Pax6. T* was already expressed in some of the day 4 *Mtf2* null cells, as compared to day 7 for wild type cells (Wilcoxon signed-rank test p value < 4 x 10^−3^; Fig. 3f). The key endodermal marker *Foxa2* was predominantly expressed in the *Mtf2* null cells but not *Jarid2* null cells (Wilcoxon signed-rank test p value < 3 x 10 ^-7^), in line with our finding that the differentiation towards early endodermal lineages was severely impacted in *Jarid2* null cells (Fig. 3g). Relatively more *Jarid2* null cells retained expression of the pluripotent marker *Klf4* at early time points (Fig. 3e), and more of them gained neuro-ectodermal *Pax6* expression over time (Fig. 3h).

In summary, our single cell transcriptome analyses of the different PRC2 mutants revealed differences in speed and direction of differentiation. Next, we sought to elucidate the underlying mechanism for this difference between the two PRC2 complexes containing either MTF2 (PRC2.1) or JARID2 (PRC2.2).

### Derepression of PRC2 targets in *Mtf2* null ES cells

We explored the gene expression differences between undifferentiated *Mtf2* and *Jarid2* null cells in more depth by bulk RNA sequencing. This uncovered a significant number of differentially expressed genes between *Mtf2* null and wild type cells, the majority of which was upregulated (Fig. 4a-b, Extended Data Fig. 4a). A large part of the variance in our dataset could be attributed to the *Mtf2* null line, whereas the wild type and *Jarid2* null cells were relatively similar (cf. undifferentiated cells, Extended Data Fig. 4b).

**Figure 4.**
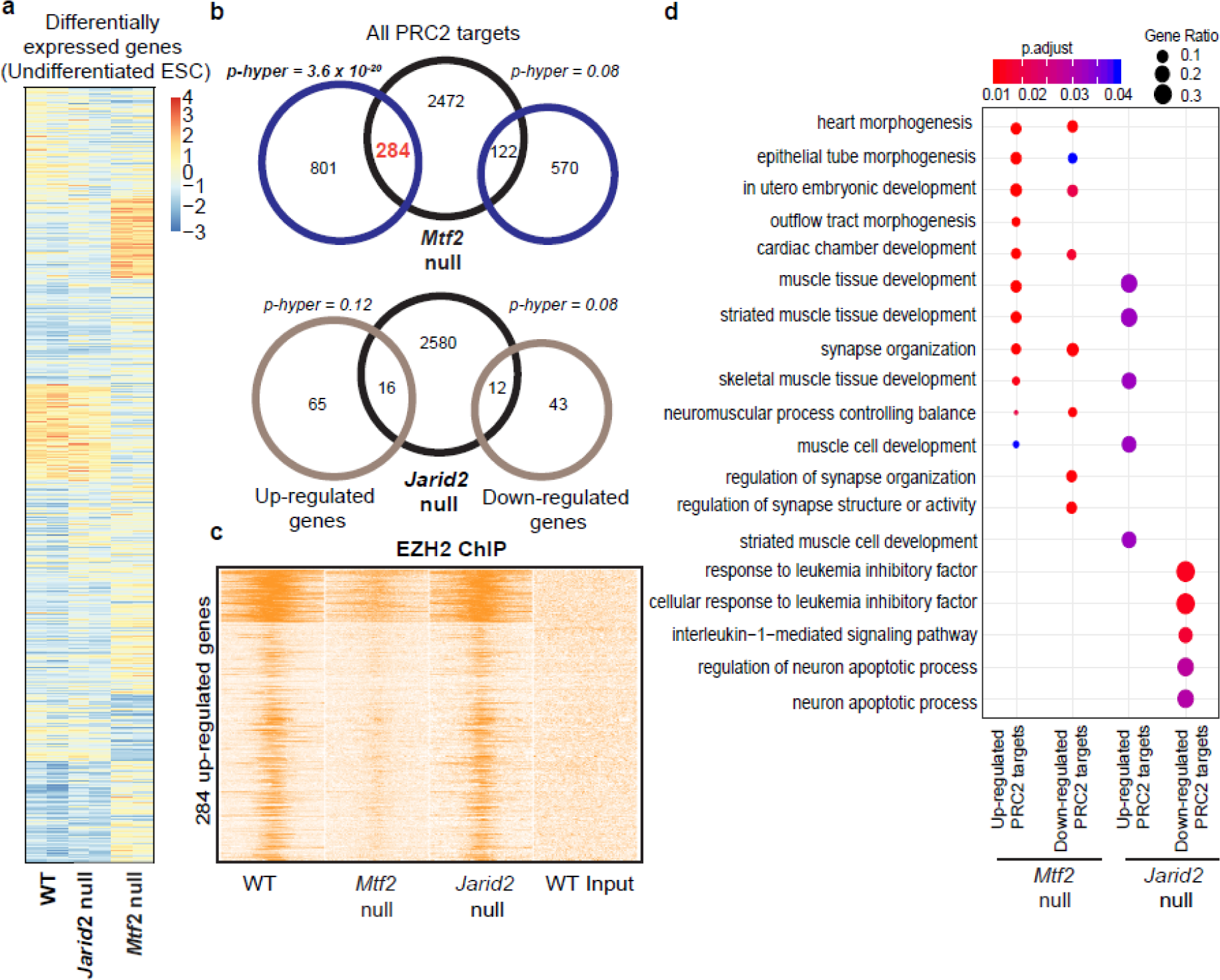
**a**, Heatmap of differentially expressed genes comparing different mutants against WT in undifferentiated mESCs. Normalized counts shown in heatmap. **b**, Venn diagram depicting overlap of number of genes which were up/downregulated in *Mtf2*^GT/GT^ and *Jarid2* null cells, with PRC2 targets (as determined from EZH2 ChIP targets; next panel). Significance of overlap calculated using hyper-geometrical test. **c**, Heatmap of EZH2 binding peaks for selected 284 target genes (from *Mtf2* null mESCs) for different genetic backgrounds. **d**, Dot plot showing the enriched Gene Ontology terms for the differentially expressed genes in different genetic backgrounds.

Next, we overlapped the up-regulated genes from both genetic backgrounds with a list of PRC2 target genes (Extended Data Table 2) which we defined by EZH2 ChIP-sequencing^13^.The up-regulated genes in *Mtf2* null cells overlapped significantly with EZH2-bound genes (284 genes, hypergeometric p value < 3.6 x 10^−20^; Fig. 4b), whereas the down-regulated genes did not show a significant overlap (Fig. 4b). This is concordant with derepression of PRC2 targets in the absence of *Mtf2*. We detected very few differentially expressed genes in *Jarid2* null cells (n=136), and neither upregulated nor downregulated genes in *Jarid2* null cells overlapped significantly with EZH2-bound genes (Fig. 4b, Extended Data Figure 4a). There is a small overlap between the genes upregulated in the absence of MTF2 and JARID2 (n=47; Extended Data Fig. 4c). We examined the levels of EZH2 recruitment of the 284 genes upregulated in *Mtf2* null cells, and observed a relatively strong depletion of EZH2 peak signals in the promoter regions of those genes in *Mtf2* null cells, whereas EZH2 recruitment to these genes was considerably less affected in *Jarid2* null cells (Fig. 4c). The small number of up-regulated genes in *Jarid2* null cells are weakly associated with specific gene ontology terms, the relevance of which, however, is not immediately apparent (Fig. 4d). The GO terms associated with LIF response are based on the down-regulation of just 4 genes in *Jarid2* null cells, one of which is *Foxd3*, which is critical for exit of pluripotency and mesendoderm differentiation ^32,33^. By contrast, the 284 up-regulated Polycomb target genes in *Mtf2* null cells were enriched for mesodermal and ectodermal tissue development Gene Ontology terms such as heart morphogenesis, muscle development and synapse organization (Fig. 4d), terms that correspond well to cell clusters with an overrepresentation of *Mtf2* null cells in the EB differentiation experiments.

Together, these analyses underscore the importance of *Mtf2* relative to that of *Jarid2* in PRC2-dependent transcriptional repression of developmental genes. The loss of *Jarid2* leads to a smaller reduction of EZH2 binding and a much less profound derepression of gene expression compared to the loss of *Mtf2*. To further understand how the up-regulated PRC2 target genes in the *Mtf2* null cells affect lineage-specific gene expression, we explored the profiles of permissive H3K4me3 and repressive H3K27me3 marks at the MTF2-dependent PRC2 target genes in mESCs.

### Key PRC2-repressed lineage transcription factors are poised for activation

Lineage commitment is associated with activation of genes specific to that lineage and repressing genes of other lineages. We, therefore, hypothesized that the primed exit of pluripotency in *Mtf2* null cells is linked to the derepression of differentiation genes. First, we analyzed the levels of repressive H3K27me3 mark at key lineage transcription factor loci, like *Sox11, Foxa2* and *Gata6* in both wild type and *Mtf2* null cells. The levels of H3K27me3 were reduced upon the loss of *Mtf2*, as expected. When we examined the permissive mark H3K4me3 in the same regions of the genes, we noticed a concomitant increase in the mark, raising the possibility that these genes were bivalent in nature and poised for transcriptional activation (Fig. 5a). A similar increase in H3K4me3 in *Mtf2* null cells is observed globally for EZH2-bound genes (Fig. 5b). Intriguingly, the 284 MTF2-dependent PRC2 target genes already had lower levels of promoter H3K27me3 as compared to the rest of the PRC2 target genes, even in wild type cells (Kolmogorov-Smirnov p-value < 5 x 10^−5^). Upon the loss of *Mtf2*, their promoter H3K27me3 levels were reduced and generally lower than the levels observed at all PRC2 targets (Fig. 5b; Kolmogorov-Smirnov p-value < 3 x 10^−13^). Moreover, the levels of H3K4me3 at these genes were higher than other PRC2 targets in wild type cells, and also increased more in *Mtf2* null cells (Kolmogorov-Smirnov p-value < 2.2 x 10^−16^), indicating that these bivalent genes were poised for activation and that H3K27me3 levels kept their expression low in undifferentiated ES cells. Previously we found that MTF2 and JARID2 mutually stabilize their binding, which in part, happens through EED binding to H3K27me3^13,34^. We noticed that overall PRC2 binding (EZH2) is rather similar and that the mutual destabilization of MTF2 and JARID2 binding is more severe for the 284 genes compared to all PRC2 targets (Fig. 5b, Extended Data Fig. 5a).

**Figure 5.**
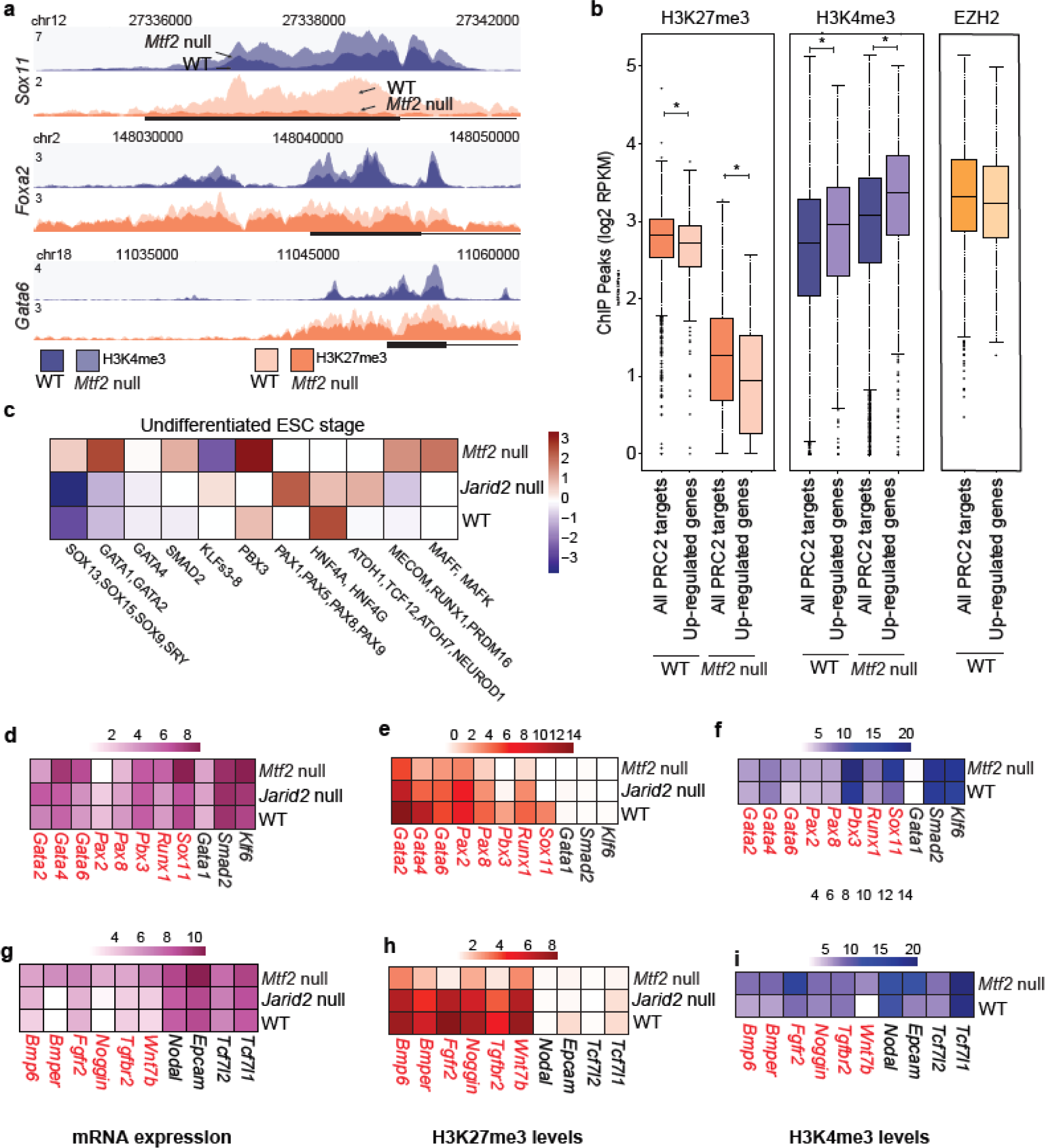
**a**, ChIP peak profiles of histone mark H3K27me3 and H3K4me3 for selected lineage transcription factors for WT and *Mtf2*^GT/GT^ cells at pluripotent stage. ChIP profiles were RPKM normalized and scaled between two merged profiles per histone ChIP. **b**, box plots depicting the log2 normalized RPKM values of promoter H3K27me3, H3K4me3 and EZH2 for all PRC2 targets and the 284 up-regulated genes in *Mtf2*^GT/GT^ cells. Asterisk (*) represents a Kolmogorov-Smirnov p-value < 0.001. **c**, Heatmap showing transcription factor motifs are associated with changes in transcript levels (Methods). Positively and negatively correlating motifs are shown in -log10(p-value) and log10(p-value) respectively. **d – f**, Heatmaps showing the mRNA expression (normalized counts), H3K27me3 and H3K4me3 levels for a set of transcription factors predicted in **c**. Genes in red are PRC2 targets, and genes in black are not. **g – i**, Heatmaps of similar datasets as **d – f** but for list of signaling factors identified from *Mtf2* null DEGs list.

This is concordant with a role for transcriptional activation in reduced H3K27 methylation at these targets, as H3K4me3 and H3K27me3 are mutually exclusive on the same histone tail. MTF2 recruitment is very similar for the 284 genes compared to all PRC2 targets in wild type cells (Extended Data Fig. 5b), and also the predicted DNA shape characteristics associated with MTF2-bound sequences^13^ are indistinguishable (Extended Data Fig. 5b). Together, these data suggest that the 284 genes are repressed by PRC2 but are poised for activation, and that PRC2 binding is destabilized by reduced H3K27 methylation, leading to derepression upon the loss of MTF2.

We analyzed transcription factor motifs to uncover gene-regulatory differences in *Mtf2* null cells versus all PRC2 targets. We regressed the motifs in the promoters of the up-regulated PRC2 targets against the variance in gene expression between different genetic backgrounds. Positive and negative contributions in the regression can be thought of as “motif activity” in the promoter regions in relation to the differences in gene expression between different lines (Fig. 5c). Among the motifs with top motif activity scores in *Mtf2* null cells were *Gata1/Gata2, Pbx3* and *Smad2*, which are known to be crucial for the exit of pluripotency and differentiation towards primitive streak and lateral plate mesoderm progenitors.

Next, we wondered if the mRNA expression of these transcription factors correlated with their predicted motif activity (Fig. 5d). We observed that the mRNA levels of some of these genes, like *Pbx3* and *Gata2*, were not correspondingly higher in the *Mtf2* null cells; these genes had similar expression and promoter H3K4me3 levels in wild type and *Mtf2* null cells, even though H3K27me3 levels at their promoters were reduced in *Mtf2* null cells (Fig. 5d-f). This suggests that, while they themselves were not more strongly expressed upon reduced Polycomb repression, their activity did contribute to stronger activation of genes that were derepressed in the absence of MTF2. Other Polycomb targets such as *Sox11, Runx1* and *Gata6* demonstrated a “poised” state of activation, with lower H3K27me3 and higher H3K4me3 levels in the *Mtf2* null cells (Fig. 5e & f). The non-Polycomb targets, like *Klf6* and *Smad2*, were expressed highly, and contained predominantly high levels of promoter H3K4me3 marks and barely any H3K27me3 signal.

In addition to transcription factors, signaling factors may also influence the exit of pluripotency. We found that many genes involved in signaling pathways such as BMP, TGFβ and Wnt signaling, were up-regulated genes in *Mtf2* null cells. Many of these signaling genes also exhibit bivalent promoter marking, and some, for example *Wnt7b, Nodal*, and *Noggin*, were more abundantly expressed in *Mtf2* null cells. Upon the loss of *Mtf2*, their promoter H3K27me3 was reduced and H3K4me3 levels were increased in conjunction (Fig. 5g-h). Together, our findings revealed that MTF2-mediated recruitment of PRC2 contributes to an epigenomic balance, the loss of which resulted in a stark de-repression of lineage transcription factor and signaling genes, and a concomitant increase in activation marks that primes embryonic stem cells towards lineage differentiation.

### Faster progression of *Mtf2* null cells to early germ layer progenitors during directed differentiation

Finally, we sought to validate our findings thus far using an *in vitro* directed differentiation system. *In vivo*, the primitive streak gives rise to mesoderm and definitive endoderm via exposure to a coordinated dose of BMP, TGFβ and Wnt signals^35–38^. We induced the formation of the primitive streak cells from mESCs over a duration of 24 hours and then bifurcated the differentiation based on activation and repression of signals which drive mesoderm and endoderm differentiation, respectively (Fig. 6a).

**Figure 6.**
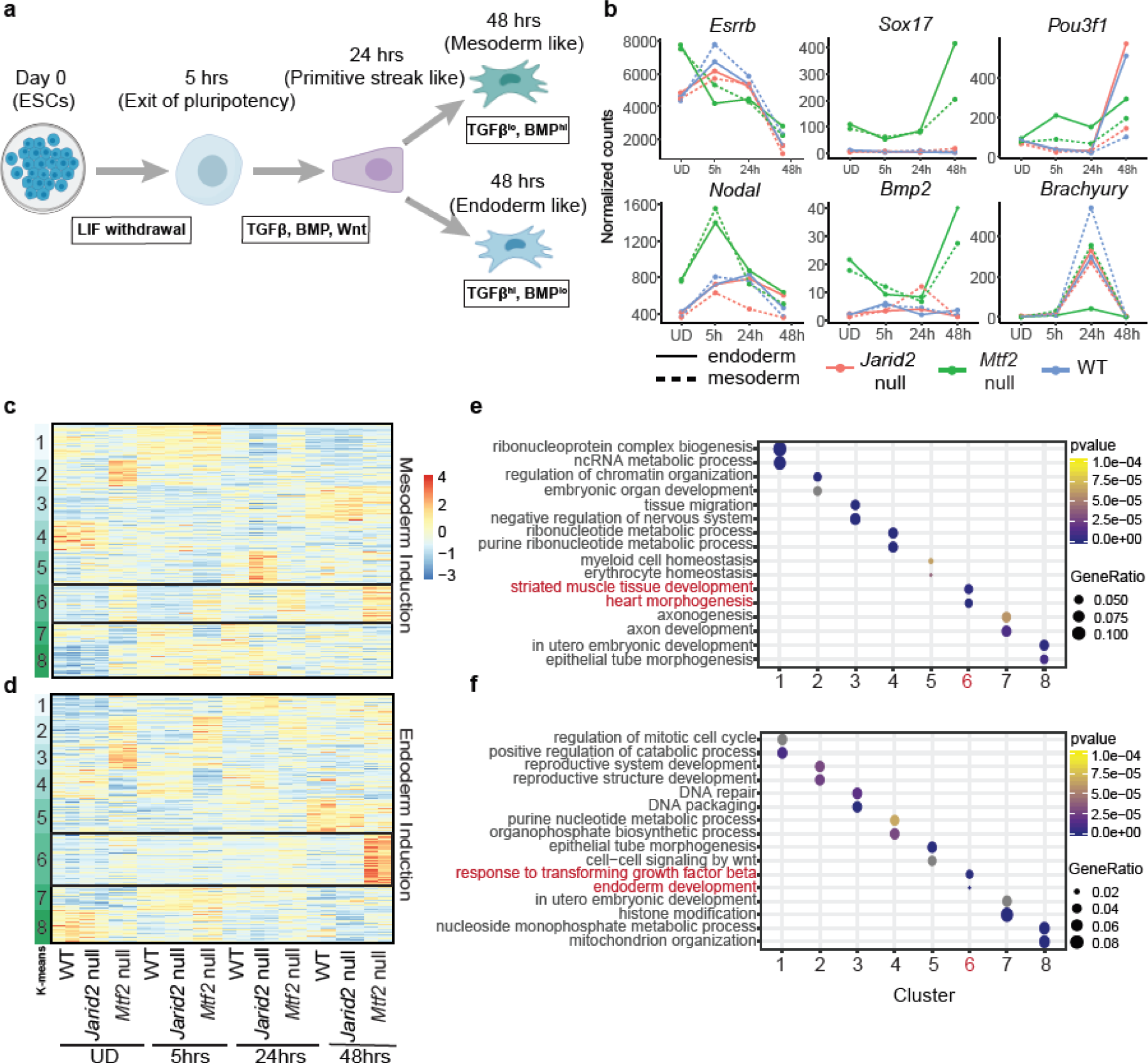
**a**, Schematic of directed differentiation for monolayer cells from different genetic backgrounds (WT, *Jarid2* null and *Mtf2* null). **b**, Line plots showing the expression (normalized counts from RNA-seq) of selected temporally regulated genes during early lineage specification processes for different genetic backgrounds and differentiation directions. **c – d.** Heatmap of differentially expressed genes across all timepoints and between genetic backgrounds. Data was k-means clustered and the normalized counts were shown. **e – f**, Dot plots showing the enrichment of biological pathways for each cluster in c – d, selected by p-value and gene-ratios of the terms. Top selected pathways were picked for each cluster and shown here.

Cells from different time points (Fig. 6a) were harvested and their bulk-transcriptomes were analyzed. The expression of certain key regulators of these early differentiation stages like *Nodal, Brachyury, Bmp2* were indeed progressively elevated, at least in the wild type cells during these early time points (Fig. 6b, Extended Data Fig. 6).

Next, we compared the differential expression of genes across different time points of *Mtf2* null and *Jarid2* null cells with the wild type cells. We performed k-means clustering to classify the differentially expressed genes and annotated each of the clusters using Gene Ontology (Fig. 6c, e). Indeed, we observed that the loss of *Mtf2* resulted in upregulation of a group of mesoderm-related genes (Fig. 6c, cluster 6) already at the undifferentiated stage. As differentiation goes on, this group of mesoderm-expressed genes became more prominently expressed, up to 48 hours of differentiation. This progressive trend was not observed for the *Jarid2* null cells. Instead, the pattern of expression for *Jarid2* null cells was remarkably similar to wild type cells.

We observed a similar trend for the endoderm lineage: the activation of endoderm genes was elevated in *Mtf2* null cells from the start (Fig. 6 d, f), whereas gene expression in *Jarid2* null cells was similar to that of wild type cells. It is also noteworthy to point out that non-endoderm gene expression in undifferentiated *Mtf2* null cells (Fig. 6d, for example cluster 2 and 3), was rapidly curbed during directed differentiation when exogenous factors directing endodermal differentiation were provided. Therefore, while undifferentiated *Mtf2* null cells exhibit multi-lineage derepression of gene expression, these data show that lineage-specific gene expression is successfully established upon lineage commitment in different directions.

## Discussion

Seminal work done on *Mtf2* (PRC2.1) and *Jarid2* (PRC2.2) has highlighted their respective importance in facilitating PRC2 binding to specific genomic loci to compact chromatin^13,14,21,39–42^. In the current study, we explored the roles of MTF2 during the exit of pluripotency and discovered that the loss of *Mtf2* alleviated the threshold for differentiation towards lineages in all germ layers, while *Jarid2* null ES cells were slower to differentiate and subsequently were re-directed predominantly towards ectodermal cell fates. Importantly, we identified an array of lineage transcription factors and signaling modulators that were upregulated upon the loss of *Mtf2* and that were inherently poised for activation. Upon the loss of *Mtf2*, promoter H3K27me3 levels were markedly reduced, while H3K4me3 was increased.

Reports on the phenotypic developmental consequences of losing PRC2 subunits during development have been diverse. The loss of *Mtf2* has been shown to be embryonic lethal by E15.5, with observations of severe anemia and growth retardation^43–45^. On the contrary, *Jarid2* null mice exhibited early embryonic lethality (as early as E10.5)^46–48^. To study the mechanisms underlying these disparities between the loss of

PRC2 subunits, we adopted the use of embryoid bodies and directed differentiation systems for better control of early differentiation intermediates and signaling paradigms to dissect the roles of PRC2.1 and PRC2.2. Importantly, single-cell RNA sequencing also allows for untangling the heterogeneity of differentiation, and for uncovering transient developmental events which are difficult to assess in early embryos^49^. For example, we were able to capture aspects of cardiac progenitor development in our dataset (Extended Data Fig. 2a), which could be applicable to the EB field and the formation of gastruloids^50,51^.

We found that Wnt signaling was affected upon the loss of *Mtf2.* This is consistent with recent evidence that *Mtf2* null mice demonstrated impairment in definitive erythroid development, and that Wnt was regulated by MTF2 during erythropoiesis^25^. We also found other signaling pathways (e.g. FGF, BMP and TGFβ) to be deregulated when *Mtf2* is mutated. These pathways were not deregulated in the *Jarid2* null cells, except for *Bmp4*, which was downregulated in *Jarid2* null cells in comparison with wild type and *Mtf2* null cells. This may potentially explain the delayed and biased EB differentiation we saw in *Jarid2* null cells, whereas it would still allow directed differentiation to proceed when exogenous signaling molecules are supplied.

Furthermore, we observed a relatively modest decrease in H3K27me3 levels upon the loss of *Jarid2*, with fewer genes that were differentially expressed. This may be due to a functional redundancy role of *Jarid2* in PRC2 function during exit of pluripotency. PRC2.1 and PRC2.2 complexes functionally interact with each other, as they both depend to some extent on positive feedback by binding of core subunit EED to H3K27me3. In the presence of EED inhibitors, PRC2 recruitment becomes much more dependent on JARID2^34^. Research done on early mESCs differentiation showed a stark increase in JARID2 stoichiometry (with respect to PRC2 core) when comparing between pluripotency and differentiated states^52,53^. Conceivably, JARID2 may play a more important role at later stages compared to undifferentiated ES cells.

In addition to signaling factors, we identified key gastrulation transcription factors with bivalent promoters that are poised for transcriptional activation. A reduction of H3K27me3 at their promoters due to the absence of MTF2, resulted in increased levels of H3K4me3. Conversely, since H3K4me3 and H3K27me3 are mutually exclusive on the same histone tail^54^, the H3K4me3 permissive mark may also limit the extent to which the promoter can be repressed by PRC2. How this epigenomic balance is regulated is a key question in the field, and our results show that MTF2-mediated targeting of PRC2 influences that balance in a way that affects the exit of pluripotency of ES cells. Of note, we also analyzed the shape of the promoters of the previously mentioned signaling and gastrulation factors but did not find a different density of DNA shape-matched GCG sequences, which can be bound by MTF2^13,39^. This suggests that indeed, reduced PRC2 recruitment may be a consequence of stronger activation rather than a reduced capacity for PRC2 recruitment *per se*. This is compatible with recent results, showing that the H3K4 methyltransferase MLL2 protects developmental genes from repression by repelling PRC2^55^, highlighting the dynamic epigenomic balance of the classical Trithorax and Polycomb systems.

In summary, our findings revealed the unique differences in differentiation speed and lineage specificity upon the loss of either of the two PRC2 subcomplexes, containing MTF2 (PRC2.1) or JARID2 (PRC2.2). During development, key signaling pathways like BMP, Wnt, FGF and TGFβ are critically regulated in a time and space-sensitive manner. The activity of these pathways causes lineage specification, but this requires coordinated responses of cells, mediated by a carefully orchestrated balance of activating and repressive cues. Our results highlight that PRC2.1 and PRC2.2 have distinct contributions in the dosage of Polycomb repression, and these differences in turn affect the balance between activation and repression of signaling pathways that are not only important in regulating the threshold for the exit of pluripotency (Fig. 7), but also during further lineage decisions during development and differentiation.

**Figure 7.**
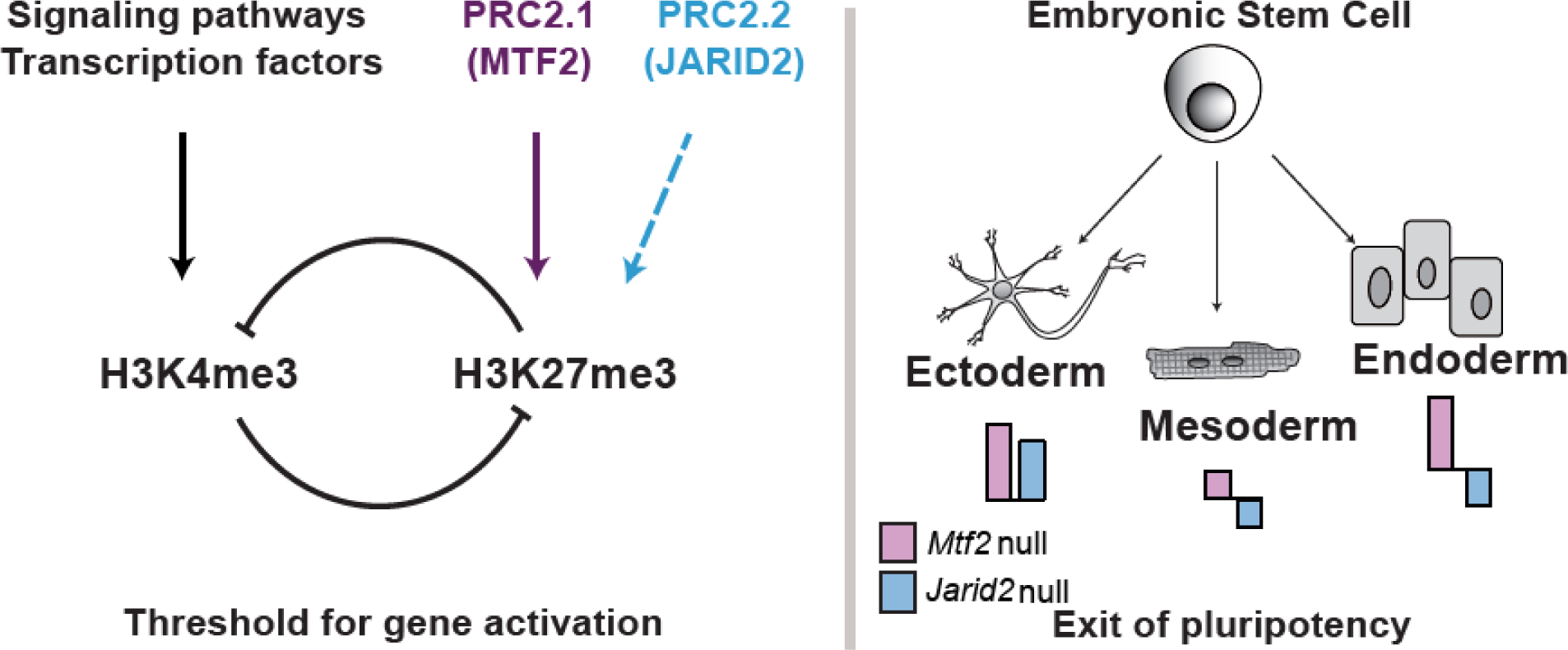
Model of the role of PRC2 in the exit of pluripotency. PRC2 contributes to a repressive threshold for activation of key regulators of differentiation (left). The different extent to which PRC2.1 and PRC2.2 contribute to H3K27me3 and repression of genes affects the exit of pluripotency and the differentiation to lineages of the three germ layers.

## Methods

### Mouse embryonic stem cell (mESC) culture

Wild-type (WT) E14 embryonic stem cells (129/Ola background) were maintained in Dulbecco’s Modified Eagle Medium (DMEM) containing 15% fetal bovine serum, 10 mM Sodium Pyruvate (Gibco), 5 μM beta - mercaptoethanol (BME; Sigma) and Leukemia Inhibitory Factor (LIF: 1000U/ml; Millipore). The E14 WT^56^, *Mtf2* null^45^, *Jarid2* null^7^ and *Eed* null^57^ cells used in this paper were maintained in serum + LIF media as described previously. Medium was refreshed once every two days.

### mESCs embryoid body (EB) differentiation

mESCs were dissociated with Accutase™ and seeded in Nunclon Sphera 6-well plates (Thermo Fisher Scientific, # 174932) at a cell density of 11,000 cells / well in Serum + LIF medium. After two days of aggregation, the embryoid bodies (EB) were let to differentiate by removal of LIF. Differentiation media was refreshed every two days by directly pipetting out spent media and adding in new media, with as little disturbance to the EBs as possible. On days of harvest, EBs are pipetted out and spun down at 400g for 5 mins, followed by dissociation with Accutase™ (37°C, 5 mins) and thereafter FACs sorted for viable cells using 7-AAD staining (Thermo Fisher Scientific, #A1310).

### mESCs monolayer differentiation

mESCs were seeded at a density of 9,000 cells / well of a 12-well cell culture plates. On the next morning, mESCs were washed (once with DMEM) and then differentiated into either anterior primitive streak (APS) by adding 100 ng/mL Activin A (Thermo Fisher Scientific, #PHC9561) + 2 µM CHIR99021(Peprotech, #2520691) + 20 ng/mL FGF2 (Thermo Fisher Scientific, # 13256029) or mid primitive streak (MPS) (30 mg/mL Activin A + 40 ng/mL BMP4 (Thermo Fisher Scientific, # PHC9533) + 6 µM CHIR99021 + 20 ng/mL FGF2) for 24 hrs. Subsequently, the cells were further differentiated into definitive endoderm with 100 ng/mL Activin A + 250nM DM3189 (Peprotech, # 1062443) and lateral mesoderm with 1 µM A-83-01(Tocris, #2939) + 30 ng/mL BMP4, respectively, for another 48hrs.

### RNA extraction and bulk RNA sequencing preparation for monolayer cells

RNA from mESCs were harvested at several time points (Day 0 undifferentiated, 5hrs after induction, 24hrs APS/MPS and 48hrs DE/LM). RNA isolation and purification were performed using the Quick – RNA™ MicroPrep kit (Zymo Research), according to the manufacturer’s protocol. Integrity of purified RNA was checked on an Agilent Bioanalyzer using the RNA 6000 Pico Kit. Intact RNA was depleted of rRNA and prepared for sequencing using the KAPA RNA HyperPrep Kit with RiboErase (Kapa Biosystems). Libraries were sequenced on the NextSeq 500 (Illumina), generating an average of 12-15million reads per sample.

### Single-cell RNA library preparation for EBs

EBs were harvested at different timepoints (Day 0 undifferentiated, 4, 7, 10 days). The cell suspension was pipetted ∼15times to prevent cell clumping and it was stained with 7-AAD.The live cells were selected for and FACs – sorted onto 384-well plates containing primers with unique molecular identifiers, according to the SORT-Seq protocol^58^. Plates were spun down (1200g, 2mins, 4°C) and ERCC spike-in mix (1:50,000) was dispensed by a Nanodrop (BioNex Inc) into each well. 150nl of the Reverse Transcription (RT) mix was similarly dispensed into each well. Thermal cycling conditions were set at 4°C 5min; 25°C 10min; 42°C 1hr; 70°C 10min. Contents from the plates were pooled together and the cDNA was purified using AmpureXP (New England BioLabs) beads. In-vitro transcription (Ambion MEGA-Script) was then carried out overnight at 16°C, with the lid set at 70°C. An exonuclease digestion step was performed thereafter for 20mins at 37°C, followed by fragmentation of the RNA samples. After a beads cleanup, the samples were subjected to library RT and amplification to tag the RNA molecules with specific and unique sample indexes (Illumina), followed by a final beads cleanup (1:0.8, reaction mix: beads) and the sample cDNA libraries were eluted with DNAse free water. Libraries were quantified using by qPCR and sequenced on the NextSeq 500 (Illumina) for 25 million reads per plate.

### Reads filtering, processing and downstream analyses pipeline

#### Single – cell RNA

Raw reads were mapped and aligned to the mouse genome GRCm38/mm10 database using the Bowtie2^59^ alignment tool. Aligned reads were indexed and the final count table was derived using HTseq^60^. R package ‘Scater’^61^ was used for sample filtering and quality check, to remove dropouts (cells with < 5 reads) and cells with too few recovered genes (<500). The data was than analysed using R package ‘Seurat’^62^ for batch, read counts and gene counts normalization. Thereafter, the filtered and normalized reads were subject to a Principal Component Analysis using the package’s inbuilt functions. The cells are then clustered using a shared nearest neighbor (SNN) modularity optimization - based clustering algorithm. Time course trajectory analyses were performed using the package ‘Monocle 2’^63^. Cluster identities were defined by Anatomy ontology^64^ and correlated with Mouse Cell Atlas^29^ (MCA) cluster data.

#### Bulk – RNA

Paired-end Illumina 75-bp sequencing files were mapped to the mouse genome GRCm38/mm10 database using the Bowtie2^59^ alignment tool. Reads were quantified using Salmon^65^ and the count tables were analysed using DEseq2^66^ (version 1.18.1), using Wald statistics(Log2 fold change > 1) for pairwise comparison and Likelihood Ratio Test statistics (FDR < 0.01) to identify statistically different expression patterns across timepoints. Gene Ontology enrichment analysis was performed with clusterProfiler^67^ (version 3.6.0). Anatomy ontology enrichment was performed MouseMine web interface^64^.

#### ChIP-seq analyses

ChIP data for undifferentiated mESCs were generated previously^13^. All fastq files were mapped using bwa (version 0.7.10-r789) and filtered using samtools (version 1.7, flag -F 1024), then normalized for depth of sequencing. Peak-calling was done using MACS2-2.7^68^ (qvalue <0.0001). Only peaks that were called in both replicates were used downstream. Heatmaps for ChIP-seq were generated using fluff^69^(v3.0.2) from bam files using read-deapth normalization. Reads Per Kilobase of transcript, per Million mapped reads (RPKM) quantification from two independent replicates were performed using scipy (v 1.1.0). GimmeMaelstrom^70^ (v 0.14.0) was used to identify the transcription factor motifs that are influencing RNA expression dynamics. DNA shape analysis was performed using the DNAshape package^71^.

#### RT – PCR

500ng of genomic RNA were used for reverse transcription into cDNA using the iScript cDNA synthesis kit (Bio-Rad, #170-8891). cDNA was diluted 10X and 2uL was added for each qPCR reaction (iQ™ SYBR Green Supermix, Bio-Rad) and the subsequently ran on the CFX96 Touch™ Real-Time PCR Detection System.

## Acknowledgements

We would like to thank Henk Stunnenberg and Peter Brazda for help with single cell RNA sequencing at early stages of the project, Eva Janssen-Megens for technical assistance, Guido van Mierlo for scientific input and discussion, and Rob Woestenenk for help with sorting cells.

## Author Contributions

CHL and GJCV designed the experiments. CHL performed all experiments and bioinformatic analyses. MP designed experiments in the early phase of the project, provided ChIP data for undifferentiated mESCs, bulk RNAseq of EB differentiation and analysis, and performed DNA shape prediction analysis. MRB assisted with the bulk RNA-seq analyses.

