## Extended Data and Supplemental Tables for "Loss of PRC2 subunits primes lineage choice during exit of pluripotency": Ext.Data.Fig.1.pdf

### Extended Data Fig.1: Gene expression in wild type and mutant embryoid bodies

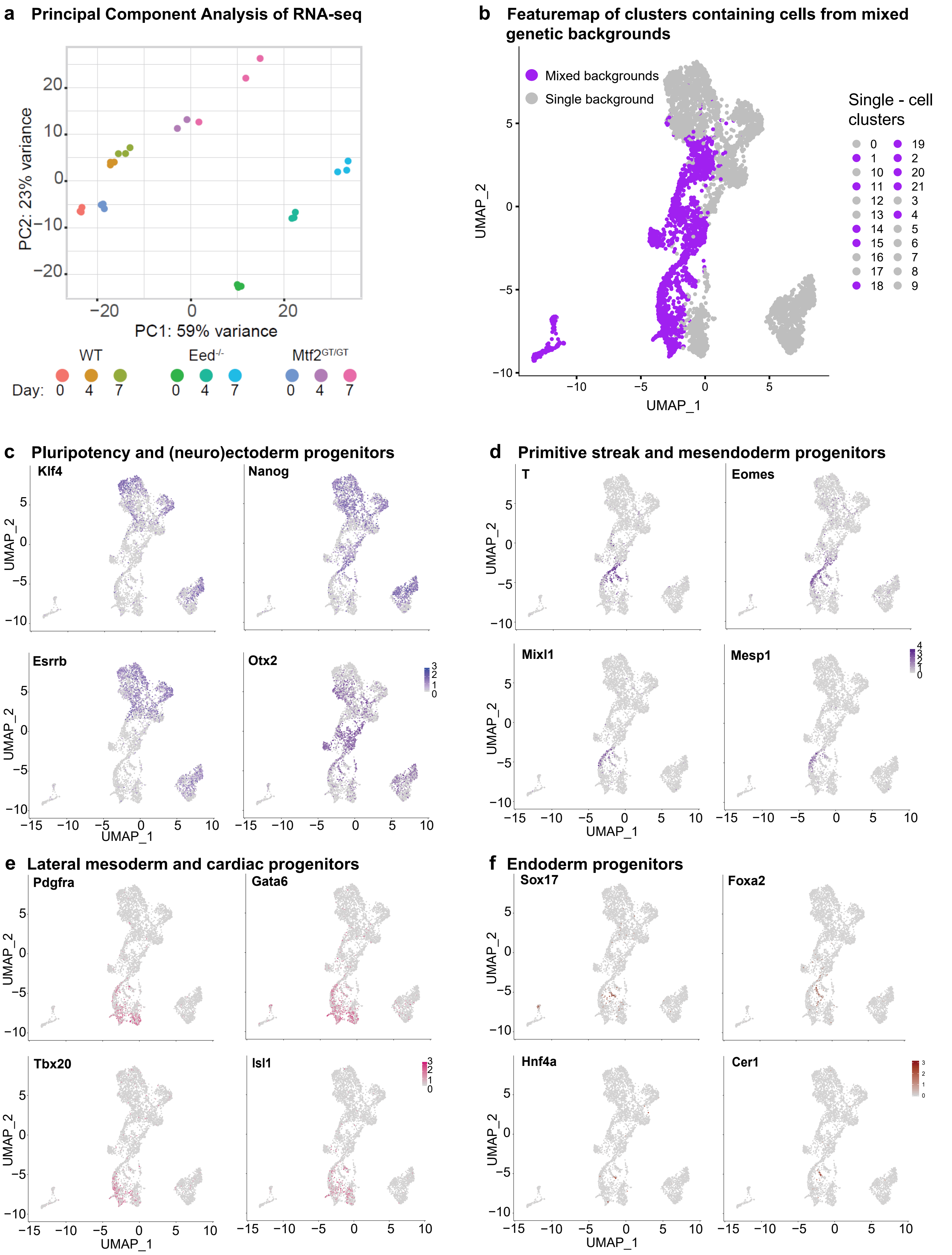


**Extended Data Fig.1**, a, Principal component analyses of bulk-transcriptomes from cells of different timepoint and backgrounds during EB differentiation. b, Feature map showing the clusters which contain mix genetic background (purple) and others which contain single background. c - f, Feature maps depicting the expression pattern and levels of selected key lineage genes from different germ layers during development.
