## Extended Data and Supplemental Tables for "Loss of PRC2 subunits primes lineage choice during exit of pluripotency": Ext.Data.Fig.2.pdf

### Extended Data Fig. 2: Cell clusters in embryoid bodies of Polycomb mutant ESCs

**a** Anatomy enrichment

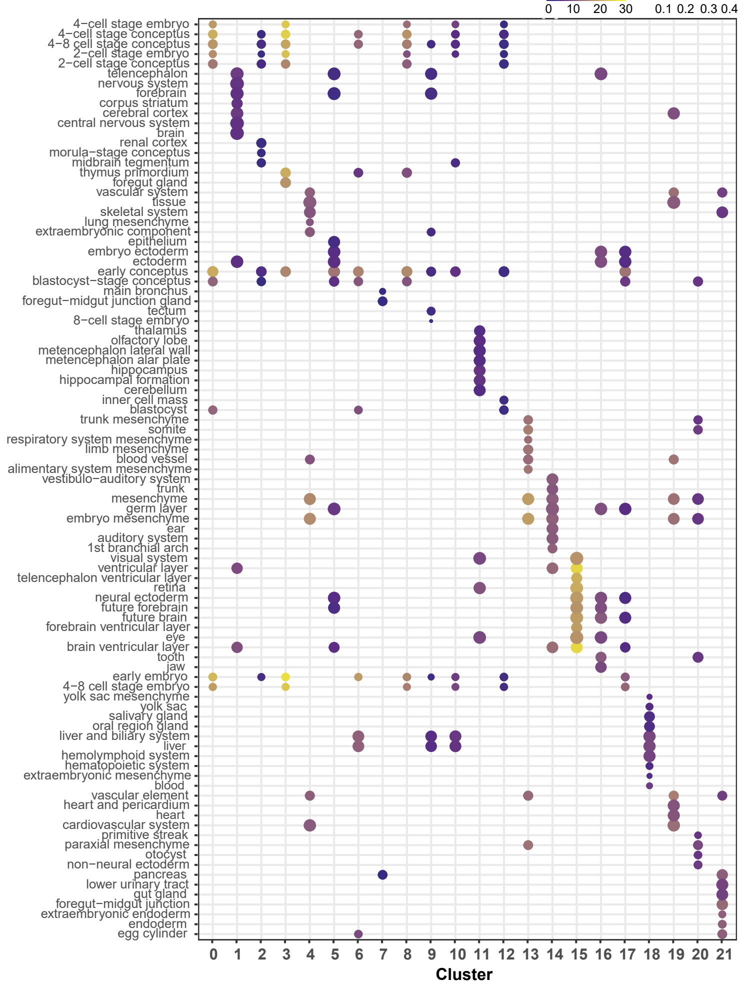


**b** Mouse Cell Atlas Enrichment

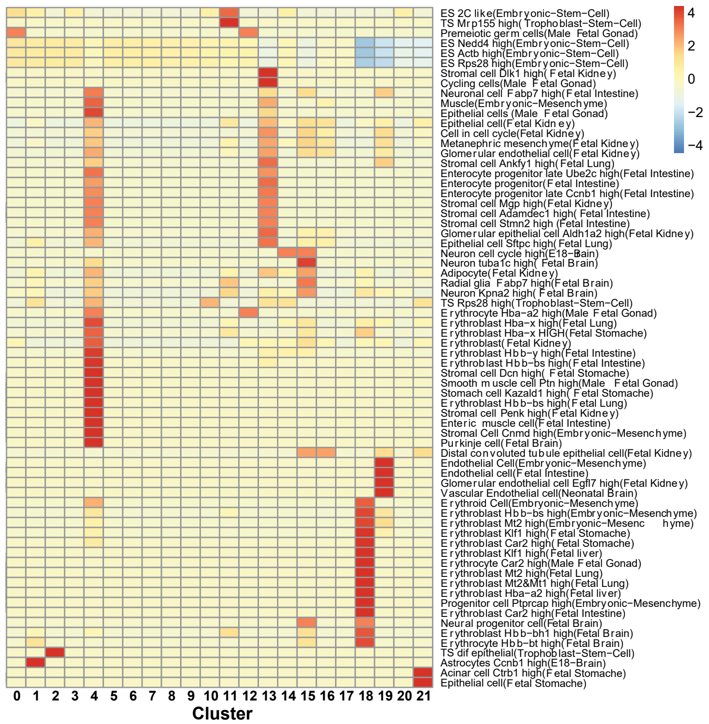


**c** Germ layer groups of single cell clusters

| Clusters | Mouse Anatomy Ontology | Broad Germ Layer Category |
|---|---|---|
| 4 | Cardiovascular System; Vascular Element | Mesodermal |
| 13 | Embryo Mesenchyme; Vascular Element | |
| 19 | Heart; Heart and Pericardium | |
| 9 | Liver; Biliary System | Endodermal |
| 10 | Liver; Biliary System | |
| 21 | Gut; Foregut - Midgut Junction; Endoderm | |
| 1 | Cerebral Cortex; Ectoderm | Ectodermal |
| 11 | Eye, Retina, Visual system | |
| 15 | Brain Ventricular Area, Future Forebrain | |
| 16 | Tooth, Jaw, Eye, Neural Ectoderm | |

**d** Single cell clusters and background information

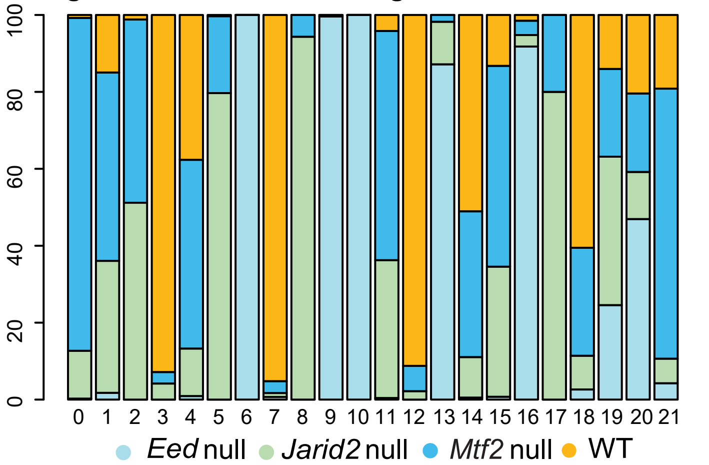


**e** Other early differentiating precursor clusters

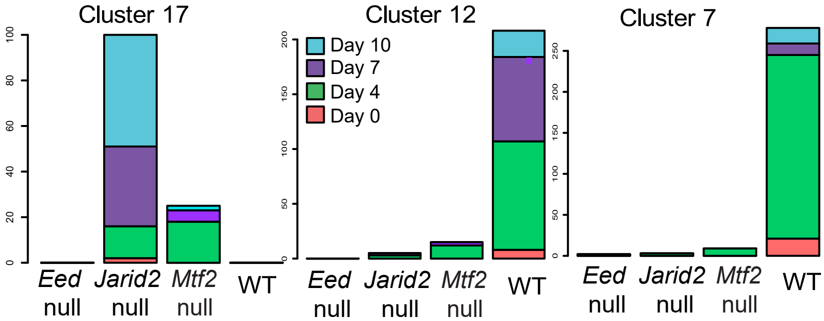


**Extended Data Fig.2 a**, Dotplot of enriched Anatomical terms representing different clusters from single-cell analyses. b, Heatmap of enrichment of single-cell clusters compared to clusters from the Mouse Cell Atlas database. c, Table of annotated clusters based on Anatomy ontology and broad germ layer classification. d, Barplot showing the proportions of cells from different genetic backgrounds (colours) in all single-cell clusters. e, Barplots of number of cells from different backgrounds in other early differentiating precursor clusters from single-cell data apart from cluster 5 shown in main panel.
