## Extended Data and Supplemental Tables for "Loss of PRC2 subunits primes lineage choice during exit of pluripotency": Ext.Data.Fig.3.pdf

**Extended Data Fig. 3: Lineage trajectory analysis confimed faster generation of cell types in *Mtf2* null cells.**

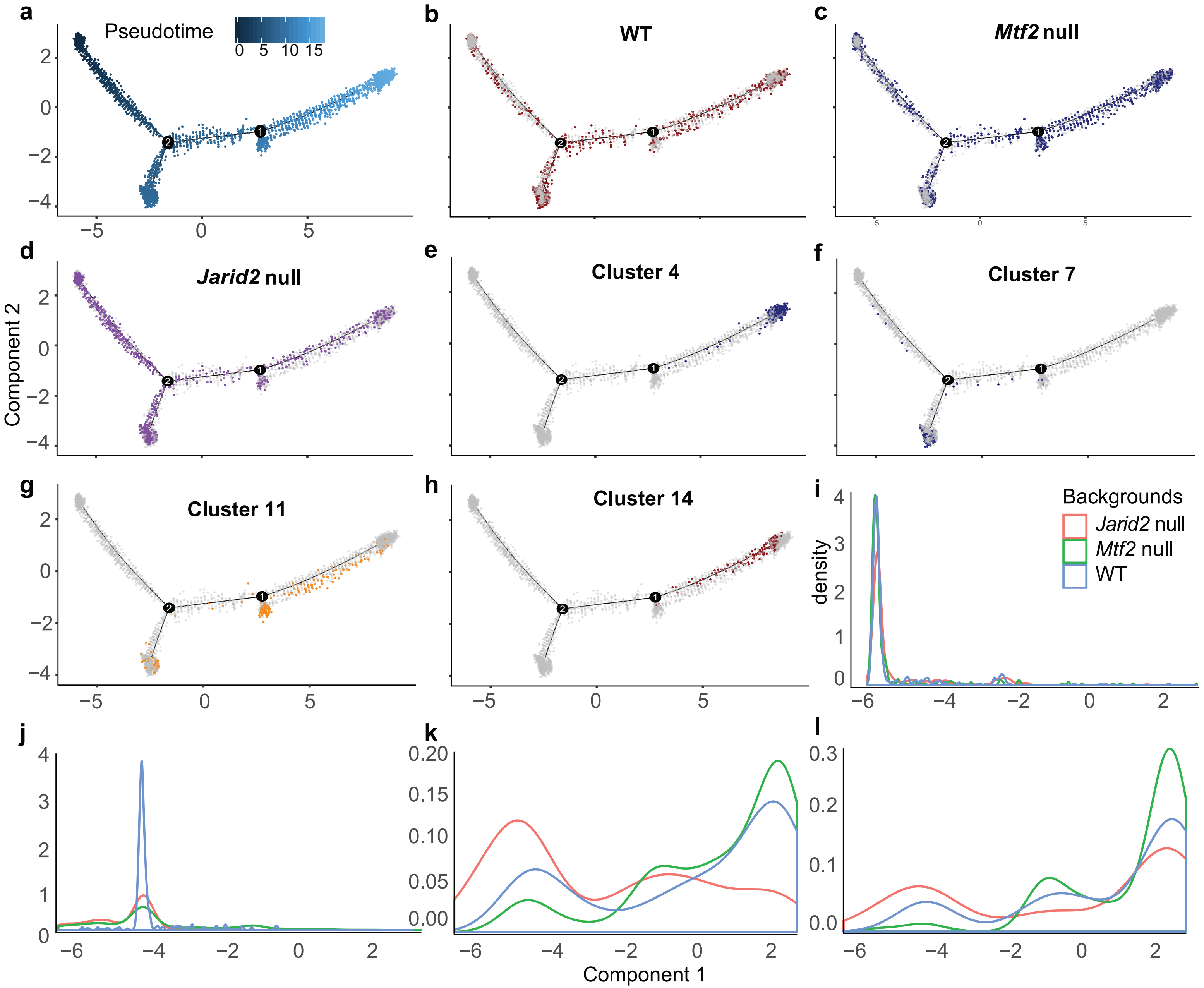


**Extended Data Fig.3** Trajectory analyses plot depicting **a,** pseudotime of individual cells **b - d,** pseudotime plots overlaid with different genetic backgrounds **e - h,** selected single cell clusters from different timepoints. **i - l,** different density plots of number of cells from different genetic backgrounds over pseudotime (x-axis)**.**
