## Extended Data and Supplemental Tables for "Loss of PRC2 subunits primes lineage choice during exit of pluripotency": Ext.Data.Fig.4.pdf

### Extended Data Fig. 4: Derepression of PRC2 targets in *Mtf2* null ES cells.

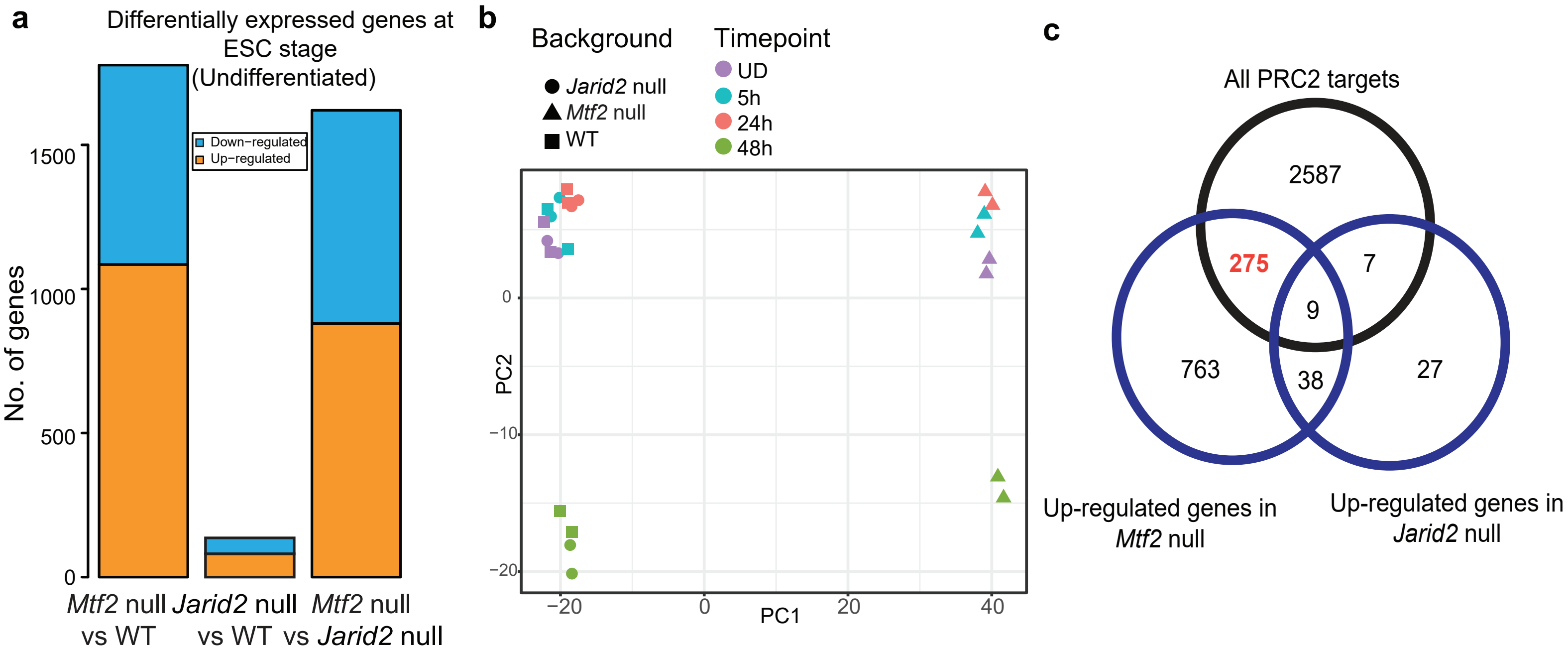


**Extended Data Fig.4 a,** Barplots showing the number of differentially expressed genes between genetic backgrounds at the undifferentiated stage in monolayer culture. **b,** PCA analysis showing variance between timepoints and background during different stages of directed differentiation on monolayer. c, Venn diagramm for overlapping up-regulated genes between *Mtf2* and *Jarid2* null.
