## Extended Data and Supplemental Tables for "Loss of PRC2 subunits primes lineage choice during exit of pluripotency": Ext.Data.Fig.5.pdf

### Extended Data Fig. 5 Key PRC2-repressed lineage transcription factors are poised for activation

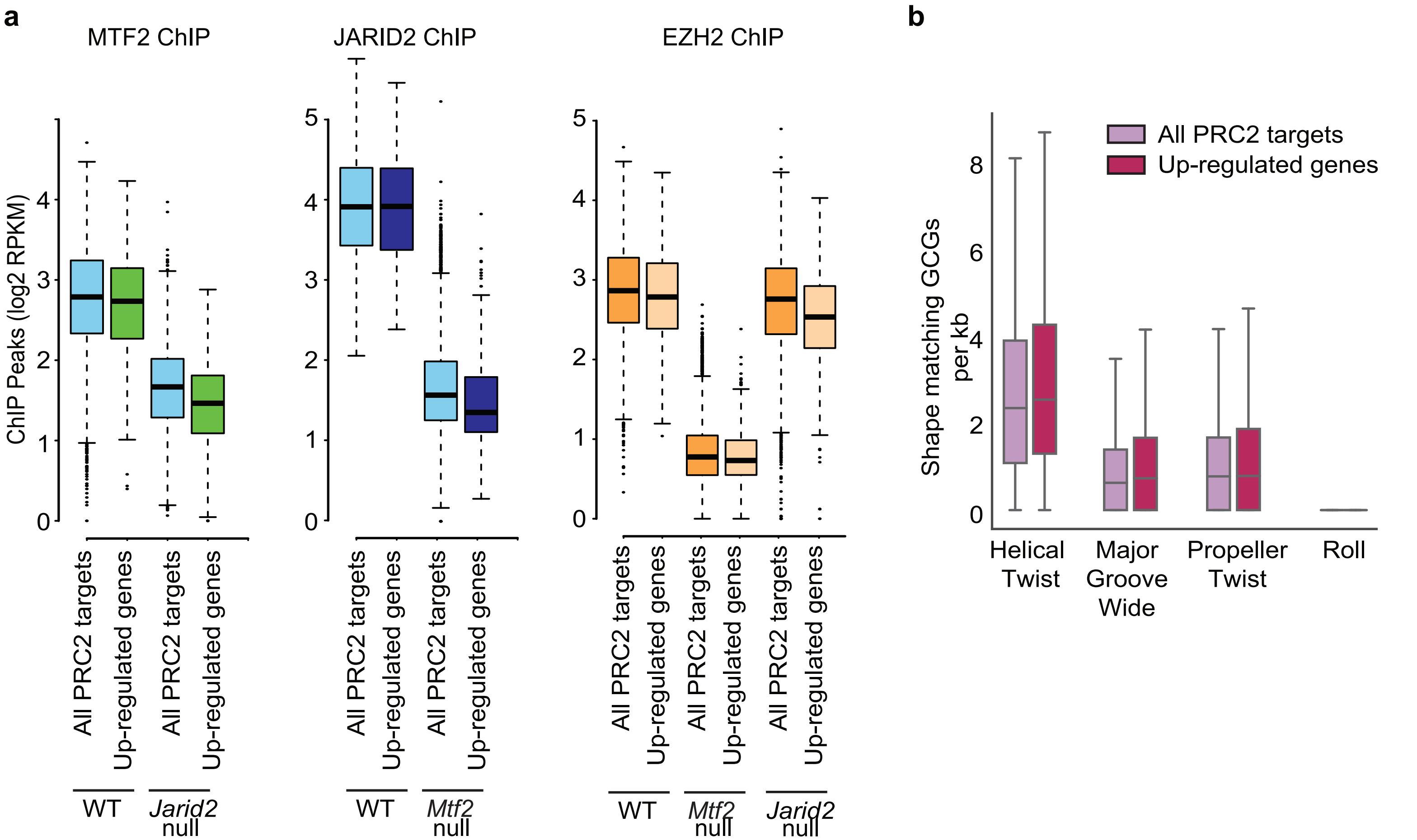


**Extended Data Fig.5 a,** Barplots showing the ChIP RPKMs for MTF2, JARID2 and EZH2 comparing between all PRC2 targets and the 284-upregulated genes in *Mtf2* null cells. **b,** Number of shape matching CpG islands in the promoters of all PRC2 targets and the upregulated genes in *Mtf2* null cells.
