## Extended Data and Supplemental Tables for "Loss of PRC2 subunits primes lineage choice during exit of pluripotency": Ext.Data.Fig.6.pdf

**Extended Data Fig. 6: Directed differentiation of *Mtf2* and *Jarid2* null cells.**

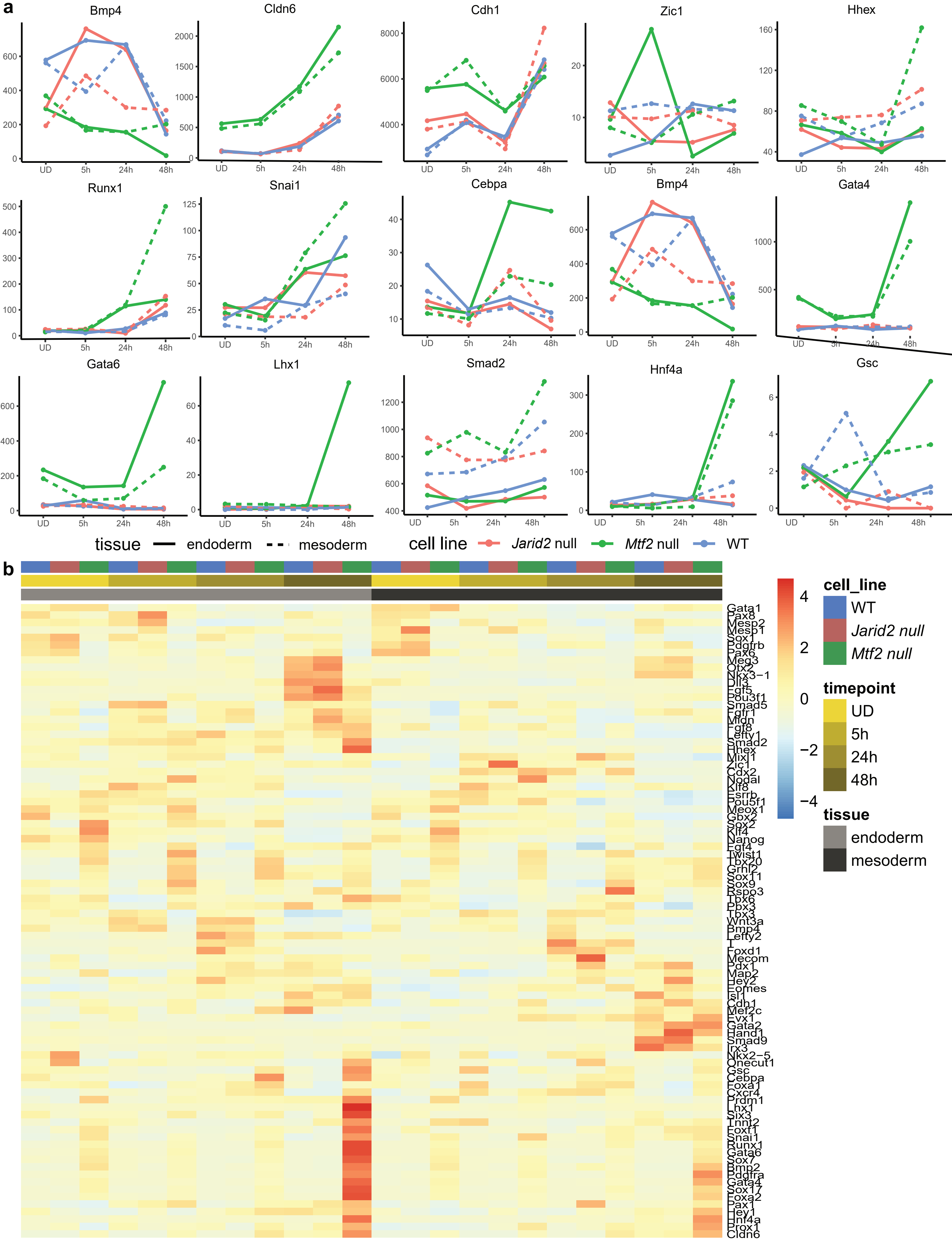


**Extended Data Fig.6 a,** Lineplots showing the expression of key early developmental factors (normalized counts from bulk-RNA seq) b, Heatmap depicting the expression differences of key mesodermal and endodermal gene markers between different differentiation experiments(mesoderma and endoderm) and across different timepoints of differentiation.
